# Exposure to traffic noise can impact multiple insect fitness components

**DOI:** 10.1101/2025.08.21.671675

**Authors:** Sarthak Saini, Anish Koner, Chinmay Gadkari, Vinayak Khodake, Sutirth Dey

**Affiliations:** Indian Institute of Science Education and Research (IISER) Pune, Dr Homi Bhabha Road, Pashan, Pune, Maharashtra, India – 411008

**Keywords:** noise pollution, *Drosophila melanogaster*, fecundity, Mating, Locomotor activity, Stress resistance

## Abstract

Noise pollution is a significant environmental stressor that affects organisms’ physiology and behavior. Due to its wide occurrence and range of component frequencies, traffic noise is a major culprit in this context. Although well-studied in vertebrate systems, the effects of traffic noise on invertebrates remain relatively less explored. This study uses *Drosophila melanogaster* as a model system to examine the impact of traffic noise on insects. Exposure to traffic noise reduced fecundity in both freshly mated and pre-mated flies, with a more pronounced effect in the former. Additionally, traffic noise increased mating latency but did not affect mating duration. The impact on locomotor activity was sex-specific: while the females remained unaffected, the male flies showed reduced activity regardless of their mating status. Desiccation resistance remained unaffected in both sexes. These findings indicate that traffic noise can decrease fitness in *Drosophila melanogaster*, which aligns with some (but not all) previous observations on other insect species. The long-term ecological implications of such noise include potential disruptions in insect-mediated ecosystem services, which can threaten the overall ecological balance.

**Summary Statement:** Traffic noise impairs fruit-fly’s fecundity, extends mating latency, and reduces male locomotor activity, suggesting that such noise can potentially reduce fitness and risk insect-driven ecosystem services.

## 1 Introduction

The post-industrialization era has experienced a significant boost in the noise levels produced due to human activities (Dimanov, 2024). The World Health Organization recognized ‘noise’ as a pollutant in 1972. Increased ambient noise (noise pollution) affects human health in multiple ways, including reduced cognitive performance, hearing loss, and adverse effects on the cardiovascular system (Babisch, 2011; Levak et al., 2008; Nassiri et al., 2013; Smith, 1989). Similarly, noise pollution can lead to increased stress hormone levels, damage parts of the hypothalamic-pituitary-adrenal (HPA) axis, and negatively impact reproduction, metabolism, heart physiology and embryonic development in animals (reviewed in Kight and Swaddle, 2011). Exposure to noise (1-7 kHz, 120 dB) induced threshold shift, apoptotic activity and reduced miRNA expression in rat cochlea (Patel et al., 2013). Permanent damage in the hair cells of the statocyst was observed in four cephalopod species when subjected to noise (50-400 Hz, 157-196 dB)(Solé et al., 2019). Geographically, noise pollution extends not only to the terrestrial regions, but also to the aquatic regions, and affects the development, physiology and behaviour of the aquatic animals (reviewed in Kunc et al., 2016). Many animals use acoustic signals to look for mates and escape predators. In such organisms, the inability to detect an essential sound cue because of the increased ambient noise (i.e., masking) can become a significant factor in determining their fitness. Due to noise, primates, birds, cetaceans, and a sciurid rodent have been shown to shift their patterns of vocal signals (reviewed in Barber et al., 2010; Erbe et al., 2018). Anthropogenic noise exposure has increased in both the noise level and the geographical scale, which induces behavioural changes in animals, e.g., deterring them from their usual foraging grounds. All these changes, in turn, could affect the evolution and species distribution across geographical regions, affecting the ecological community dynamics (reviewed in Kok et al., 2023; Slabbekoorn et al., 2018).

Amongst all sources, transportation networks have become “the most spatially extensive source of anthropogenic noise”. With an estimated 3.0-4.7 million km of new roads to be added to the Earth’s surface by the year 2050 (Meijer et al., 2018), traffic noise is expected to become an even bigger polluter in the coming years. Besides being widespread, traffic noise poses a unique challenge for an organism. Traffic noise is heterogeneous, encompassing a wide frequency range (50 Hz to 10,000 Hz (Can et al., 2010)) and intensity (noise level ranging from 82 dBA for a car, 90 dBA for a motorcycle, 103 dBA for a heavy cargo truck to 150 dBA for a turbojet airplane (Grubesa and Suhanek, 2020)), which can potentially be more detrimental to organisms than noise sources with a single frequency or intensity.

While the effects of anthropogenic noise in general, and traffic noise in particular, have been well-investigated in vertebrates, there are relatively few studies on the invertebrate systems. This is unfortunate because arthropods alone (to say nothing about the other invertebrate phyla) constitute 90% of the Earth’s biodiversity (Wilson, 1997). Due to their sheer numbers and widespread distribution, invertebrates are likely to be affected by ever-increasing noise pollution. It is known that insect-plant interactions modulate nutrient cycling in nature and increase the primary productivity of the ecosystem (Belovsky and Slade, 2000). Insects also interact with the food web (predation, microbial feeding, detritivore) and provide ecosystem services (like pollination and seed dispersal), which affect the plant and animal species composition (Weisser and Siemann, 2013; Yang and Gratton, 2014).

There is a small but growing body of work on the effects of traffic noise on insects (Bunkley et al., 2017; Cammaerts and Cammaerts, 2018; Gurule-Small, 2018; Gurule-Small and Tinghitella, 2019; Lampe et al., 2012; Rebar et al., 2022; Welsh et al., 2023), which has highlighted that traffic noise can indeed play a detrimental role for insect physiology. When field crickets (*Teleogryllus oceanicus*) are exposed to traffic noise with consistent intensity during development, adulthood, or both, relatively few fitness traits are affected (Welsh et al., 2023). However, when exposed to traffic noise with heterogeneous profiles (a large range of fluctuating intensity), the same species suffered delayed maturity (Gurule-Small and Tinghitella, 2019), reduced longevity (Gurule-Small and Tinghitella, 2019) and impaired mate location ability (Gurule-Small, 2018). Interestingly, though, all invertebrates are unlikely to be affected similarly by noise, and at least in some cases, the effects of noise can be positive or neutral. For example, it has been shown that vehicular noise can increase the dietary richness of grasshoppers (Senzaki et al., 2024). Similarly, increased anthropogenic noise levels can have positive, negative or no effects on the abundance of arthropod families (Bunkley et al., 2017). Thus, there is much context-specificity in the impact of noise on insect physiology and behaviour.

The common fruit-fly *Drosophila melanogaster* has been an important model system for understanding the effects of noise on the physiology of insects. In this species, the courtship song (or the ‘love song’) is one of the largest components of the male courtship behaviour (Lasbleiz et al., 2006). The love song is a courtship behaviour where the male vibrates its wings at a particular frequency to provide auditory courtship cues to the female. The frequency components of the love song are ∼160 to 170 Hz (sine song) and around 280 Hz (pulsed song) (Cowling and Burnet, 1981; Kyriacou and Hall, 1989). As a result, there is a possibility of ‘masking’ (Samarra et al., 2009) of courtship signals due to overlapping frequencies in the traffic noise. Exposure to loud sounds can also induce stress in *D. melanogaster* by impairing the auditory system function and changing the neural mitochondrial size (Christie et al., 2013). Specifically, acoustic trauma (“sound-induced injury to the neural mechanism of hearing” (Saltzman and Ersner, 1955) at 400 Hz induces physiological changes (increased oxidative stress, increased apoptosis and reduced mitochondrial number) as well as behavioural responses (reduced climbing and aggressive behaviour) in *Drosophila melanogaster* (Dhar et al., 2020). In *Drosophila,* Johnston’s organ (JO), present in the second antennal segment, serves the purpose of a hearing organ (Todi et al., 2004). It has been shown that noise exposure can increase the reactive oxygen species production (ROS), a marker of oxidative stress, in JO (Dhar et al., 2020). High oxidative stress can lead to temporary or permanent damage to the organ, which can manifest in terms of physiological responses in the organism.

In this study, we used laboratory populations of *Drosophila melanogaster* to investigate what happens to the life history and behaviour of flies exposed to loud traffic noise. We asked whether the mating status of the flies (pre-mated vs freshly-mated) affected their sensitivity to noise (if any) in terms of fecundity, desiccation resistance and locomotor activity. To the best of our knowledge, barring a few investigations on orthopterans, almost no studies simultaneously look at the effects of traffic noise on multiple life-history characters in the insects. Our working hypothesis, based on previous studies on crickets (Gurule-Small and Tinghitella, 2018; Gurule-Small and Tinghitella, 2019), was that noise exposure would reduce the fecundity of the flies and negatively impact other behavioural and life-history traits. We also investigated whether the adverse effects of noise on fecundity (if any) were modulated through impaired mating. This was motivated by the observation that traffic noise can potentially mask *Drosophila* songs (Samarra et al., 2009), which can potentially interfere with mating. We used a traffic noise clip with a wide frequency and intensity range to simulate an environment with traffic noise exposure for the fruit flies. We found that traffic noise exposure (during adulthood) reduces fecundity, adversely affects mating behaviour and reduces locomotor activity in males. The results of the study depict an overall fitness cost for the fruit flies inflicted by traffic noise exposure.

## 2 Methods

### 2.1 Experimental Population

The experimental flies are derived from a large (breeding population ∼2400 flies) laboratory population of *Drosophila melanogaster*, NDB_1_ (New Dey’s Baseline), whose maintenance regime has been mentioned elsewhere (Vibishan et al., 2023). NDB_1_ population is maintained under a 14-day discrete generation cycle at a temperature of 25°C in 24-hour light conditions. Eggs are collected from NDB_1_ in 37 ml plastic vials, with approximately 6 ml of standard banana jaggery medium (henceforth referred to as food) at an egg density of ∼30 eggs/vial. The day of egg collection is considered the 0^th^ day.

The adult flies start eclosing during the 8^th^ to 10^th^ day after egg collection. During the period of eclosion, the virgin flies were sex-separated every six hours under light CO_2_ anaesthesia and collected in food vials at a density of 10 flies/vial. This ensures that the flies remain virgin, as in *Drosophila*, there is no mating during the first 12 hours after eclosion (Demerec and Kaufman, 1996). At this point, we separated the flies into two groups. In the first group, both males and females were held in sex-separated vials (hence non-mated). For the second group, the flies continued to be held in the same mixed-sex vials in which they had emerged till day 12, when all the assays (except fecundity) were performed. For the fecundity assay performed on day 13, the flies in a vial were shifted to a fresh food vial on day 12. This was done to avoid the disturbance in the food medium created by the larvae hatching out of the eggs laid by the freshly eclosed adults. Thus, all the flies in the second group were expected to have mated by the time the assays were performed.

### 2.2 Sound

The flies were exposed to a sound with the characteristics of traffic noise by playing an audio clip (3 minutes and 2 seconds) on a loop during the assays (the traffic noise exposure was given only to the adult flies used in assays). The audio clip was called “City Traffic Noise” (https://audiojungle.net/item/city-traffic-noise/25418546) and was purchased from Envato Market. The noise frequencies in this clip ranged from ∼100 Hz to 10,000 Hz, and the volume of the clip was set such that the intensities ranged from ∼90 to 105 dBA, which aligned with the frequency and intensity ranges of traffic noise (Can et al., 2010; Grubesa and Suhanek, 2020). The frequency and intensity components of the noise clip were measured using a microphone connected to a spectrum analyzer application (“Audio Spectrum Analyzer Pro” by Vlad Polyanskiy on App Store) at a distance of ∼15 cm from the source speakers. The traffic noise clip was played using a pair of speakers (model: V620SILVER-“Amazon Basics USB-Powered PC speakers with dynamic sound,” (10 cm × 6.6 cm × 7.3 cm), 5.0 V, 1 A) kept at a distance of ∼15 cm from the flies.

For the mating and fecundity assay, the flies were exposed to the traffic noise in a wooden soundproof chamber (Fig. S2) measuring 53 cm × 33 cm × 40 cm. The wooden chamber was well-lit from the inside using LED strips, which emitted very little heat, and thus the temperature remained roughly similar to the ambient temperature of 24 ± 1 °C. Locomotor activity and desiccation resistance assays occurred inside the incubator (noise-exposed and control treatments in separate incubators), set at 25°C and 60% humidity.

### 2.3 Assays

All assays were performed while the treatment flies were experiencing noise. All the assays were conducted on a separate set of untested adult flies, except in the case of locomotor and desiccation assays, where the same individuals were assayed for two phenotypic traits, i.e., locomotor activity and desiccation resistance (more details in the following sections). Since the duration of the noise differed across assays, we mention them separately for each assay below. No prior data was available to estimate the variation upon noise exposure. Nor did we have any way to *a priori* predict the effect sizes of the changes. Therefore, following existing recommendations (Norman et al., 2012), we used standard sample sizes used in the *Drosophila* life-history literature for various assays (e.g., Tung et al., 2018). Moreover, there was no specific inclusion/exclusion criteria, and the flies were assigned randomly (i.e. haphazardly) to various treatments during the assays. Our assays did not require the blinding of the experimenters, since no subjective scorings were involved.

#### 2.3.1 Fecundity

The fecundity of the flies was measured on the 13^th^ day post egg-collection. The ‘fecundity tubes’ were pre-made for the experiment by cutting off the conical part of 50 ml centrifuge tubes and covering the open end with a muslin cloth. A small cup was glued to the cap of the centrifuge tube where the food was poured (’fecundity cup’). To check how different the profiles of the noise clip outside and inside the fecundity tubes were, we used a microphone to record sound outside and inside the fecundity tube when our traffic noise clip was played from outside (kept ∼15 cms from the fecundity tube). The fecundity tube was covered with muslin cloth from one end and a fecundity cup on the other. The frequency range recorded outside and inside the fecundity tubes was compared. We observed that a single layer of muslin cloth expectedly led to a slight reduction in the decibel level but no major changes in the frequency profile of the clip inside the fecundity tube (see supplementary Fig. S3). This is particularly true in the 100-300 Hz range, i.e. the frequency range of the courtship songs of the flies.

We then aspirated a male and a female fly into each fecundity tube. We used two treatments in this assay. In the “freshly-mated” treatment, both flies were unmated at the time of introduction into the fecundity tubes, whereas in the “pre-mated” treatment, both flies came from mixed-sex vials. Due to logistical reasons, the “freshly-mated” and the “pre-mated” fecundity experiments happened at different times (the freshly-mated fly fecundity assay occurred in July-August 2023, and the pre-mated fly fecundity assay was conducted in June 2024).

We assayed 35 independent replicates for each group of flies (with sound and without sound) within each treatment. The flies were exposed to noise for 12 hours inside the wooden chamber (section 2.2), and corresponding controls were kept undisturbed outside the wooden chamber during that period. Eggs laid in the fecundity cup in the 12-hour window were counted under a light microscope.

To control for any effect of lighting and environment inside and outside a wooden chamber, we conducted control fecundity experiments in which flies were kept undisturbed inside and outside the wooden chamber. We did not observe a statistically significant change in fecundity in the absence of traffic noise, irrespective of whether flies were kept inside or outside the wooden chamber (see supplementary Fig. S4).

#### 2.3.2 Mating

Since previous occurrences can change the mating behavior of flies, this trait was measured only on the ‘unmated’ flies. For this, we poured approx. 2 ml of 1.3% w/v agar (Bacteriological grade, HiMedia) into cylindrical vials (90 mm height, 25 mm diameter) to cover the base of the vial. Once the agar had solidified, we introduced one mating pair (one male and one female) into a vial. A sponge plug (∼1.5 cm in thickness) was used to plug the open end of the vial. The thickness of the sponge plug was standardised by confirming that the frequency and intensity of sound inside and outside of the vial were the same (as mentioned in section 2.3.1 for fecundity tubes). The plug was inserted into the vial such that there was a gap of 1 inch between the surface of the plug and the agar. Thus, the “mating space” available to each pair had a diameter of 25 mm and a height of 1 inch. The control setup was kept undisturbed outside, and the treatment setup was exposed to the noise inside the wooden chamber. Control and treatment groups consisted of 40 mating pairs each.

The time when a mating pair was put into the mating space was called the “setup time” for that particular mating pair. We recorded 2-hour mating videos for each treatment and manually scored those videos to note down two quantities. We measured the “mating latency” as the time taken by the flies to initiate copulation (male stably mounts on the female as visible with the naked eye of the observer), i.e.

> *mating latency = “time point at which copulation started” – “setup time”*

Similarly, the “mating duration” was quantified as:

> *mating duration = “time point at which copulation ended” – “time point at which copulation started.”*

In all the replicates, the mating latency and mating duration were recorded only for the first mating experienced by the mating pair. Once the first mating was over, the mating pair was discarded.

#### 2.3.3 Locomotor Activity

Locomotor activity was measured using the TriKinetics *Drosophila* Activity Monitor (DAM2 system, Trikinetics Inc., PO Box 325, Princeton, MA 01541 USA). This measurement was performed on both the unmated and the pre-mated flies. In the DAM system, individual flies are put in thin glass tubes (henceforth, DAM tubes) with a diameter of 5 mm and a length of 65 mm. An infrared beam runs through the middle of the DAM tube. Whenever the fly walks across the infrared beam, DAM records it as “one activity bout.”

Individual flies were gently aspirated into the DAM tubes and plugged using small cotton plugs on both ends. 32 males and 32 females in each treatment (pre-mated and unmated) were exposed to traffic noise. The control setup consisted of the same replicate size without noise exposure.

Locomotor activity (and desiccation resistance) assays were done simultaneously on both ‘pre-mated’ and ‘unmated flies’ in the same environment. We exposed 32 males and 32 females per treatment (i.e., 64 flies per treatment with an equal number of males and females) to traffic noise. The control group also consisted of 64 pre-mated and 64 unmated flies with a 1:1 sex ratio, but did not experience any exposure to traffic noise.

The locomotor activity was measured using the number of “activity bouts” recorded by DAM during the 6-hour window.

#### 2.3.4 Desiccation Resistance

Desiccation resistance was measured using the same setup to assess the locomotor activity (and therefore had the same replicate sizes) (ref. section 2.3.3). Conducting the desiccation resistance assay was possible in the same setup as the locomotor activity assay because there was neither food nor water inside the locomotor tubes. When there was no activity in the tube for at least 12 hours, the time of the last recorded activity was considered a proxy of desiccation resistance. The replicate size of the treatment and control groups was 32 per sex. Here, the time to death was recorded as the difference between when the flies were introduced into the tube and when the last activity bout was recorded. The observation for any tube was stopped if there was no activity for at least 12 hours. This assay was also performed on both the unmated and the pre-mated flies.

### 2.4 Analysis

Separate two-way Analysis of Variance (ANOVAs) were performed to analyze the effects of mating [fixed factor with two levels: unmated (freshly-mated in case of fecundity assay) and pre-mated) and noise exposure (fixed factor with two levels: with and without noise) on various assayed fitness traits. The only exception was mating behaviour, which was assayed only on unmated flies, and therefore, 1-way ANOVA was performed. The two-way ANOVAs allowed us to test for the interactions between mating status and noise exposure. However, it also meant that whenever the interaction effects were non-significant (which happened in every single ANOVA in this study!), we could not compare the unmated or the pre-mated populations regarding the impact of noise. To ameliorate this issue, following each two-way ANOVA, we performed Welch’s t-tests to compare the means of flies exposed to traffic noise (“With Sound”) with the corresponding controls that were not exposed to traffic noise (“Without Sound”) separately for the unmated (freshly-mated in case of fecundity assay) and the pre-mated regimes. We then used the Holm-Šídák step-down procedure (Abdi, 2010) to correct the p-value generated from the t-test for the inflation of the family-wise error rate. We also calculated Cohen’s *d* as a measure of effect sizes, which were interpreted as follows: “small” (*d* < 0.*5*), “medium” (0.5 > *d* > 0.8) and “large” (*d* > 0.8) (Cohen, 2013).

The fecundity assays on the unmated and pre-mated flies were performed at different times; therefore, comparing them using two-way ANOVA is not appropriate. Hence, the two-way ANOVAs were not performed on the fecundity data.

Since our sample sizes for all assays were >30, and we only compared the sample means across treatments, the central limit theorem assures that the group means are normally distributed. For every statistical test, the sample sizes of the with-sound and the without-sound treatments were exactly equal. Under these conditions, t-tests and ANOVA become very robust to departures from homoscedasticity (Quinn and Keough, 2002). Since the data for all our traits were generally symmetrically distributed (See Figs. 1-4), with no major differences in variance, we did not conduct formal tests for normality or homoscedasticity (Quinn and Keough, 2002). All statistical analysis were performed on STATISTICA (v7 Statsoft Inc, Tulsa Oklahoma, USA).

## 3 Results

### 3.1 Traffic noise reduces fecundity of freshly-mated as well as pre-mated flies

We performed a Welch’s t-test with Holm-Šídák correction (for two comparisons) to assess the effect of traffic noise exposure on the unmated and pre-mated flies separately. When pairs of unmated flies of the opposite sex were put together for 12 hours, we observed a significant decrease in the number of eggs laid by the group exposed to traffic noise compared to the control group (t_59.66_ = 4.813, *p* = 2.2 × 10^−5^, *d* = 1.15, large) (Fig. 1A). To check the reproducibility of these results, we repeated the experiment twice on independent sets of freshly-mated flies. In both the repeats, we observed a significant reduction in the fecundity of the flies exposed to traffic noise (t_55.35_ = 4.545, *p* = 3.03 × 10^−5^, *d* = 1.086, large) (Figure S1A) and (t_67.90_ = 3.754, *p* = 0.0004, *d* = 0.897, large) (Fig S1B).

**Figure 1:**
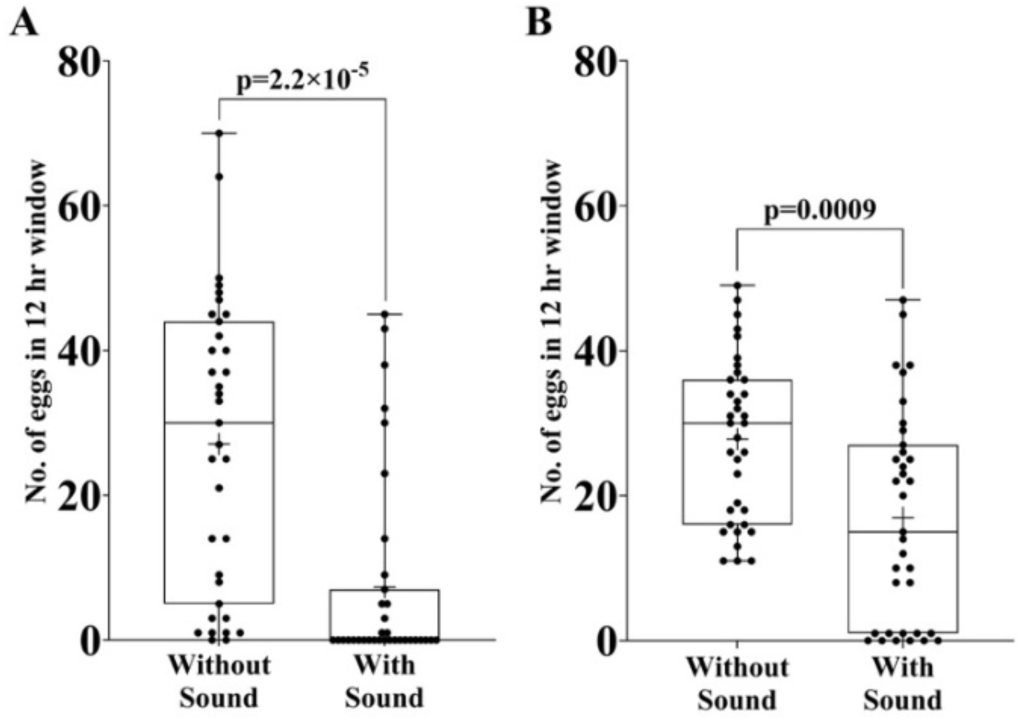
Effect of traffic noise on fecundity. A. Freshly-mated B. Pre-mated. n=35 pairs per treatment per experiment. Each pair consisted of a male and a female sampled independently from a large (∼2400 individuals) outbred lab-maintained population. The experiment on freshly-mated flies was repeated twice on the independent set of flies (data given in supplementary information). Data points are shown by the jitter on the plot. The top and bottom edges of the box denote the 75^th^ percentile and the 25^th^ percentile, respectively. The solid line between the edges represents the median. The whiskers denote the minimum and maximum data points. ‘+’ denotes the mean. Welch’s t-test (two-tailed) with a 95% confidence interval was performed to compare the means of the two treatments (i.e. with Sound and without Sound). P-values have been adjusted using the Holm-Šídák step-down procedure (for two comparisons). The data clearly shows that exposure to traffic noise reduced the fecundity of freshly-mated and pre-mated flies.

When the same experiment was performed with the flies that were expected to have mated before the experiment was conducted, we again observed a reduction in fecundity of the group exposed to the traffic noise compared to the control group (t_63.70_ = 3.479, *p* = 0.0009, *d* = 0.832, large) (Fig. 1B). However, compared to their respective controls, the magnitude of the reduction was less for the pre-mated flies (39.05 % reduction in mean relative to the respective control group) than for the un-mated flies (73.00 % reduction in mean relative to the respective control group) (Fig 1A and 1B). This led us to hypothesize that perhaps the reduction in fecundity was due to issues with mating under traffic noise.

### 3.2 Traffic noise affects the mating latency but not the mating duration

Compared to the control group, the mean mating latency of the flies exposed to traffic noise increased significantly by 71.4 % (t_63.69_ = 3.070, p = 0.0031, d = 0.733, medium) (Fig. 2A). We did not observe a significant difference in the mating duration of the group exposed to the traffic noise compared to the control group (t_69.96_ = 0.9363, p = 0.3523, d = 0.218, small) (Fig 2B). This implies that although noise interfered with mating initiation, it did not affect the duration once copulation was initiated.

**Figure 2:**
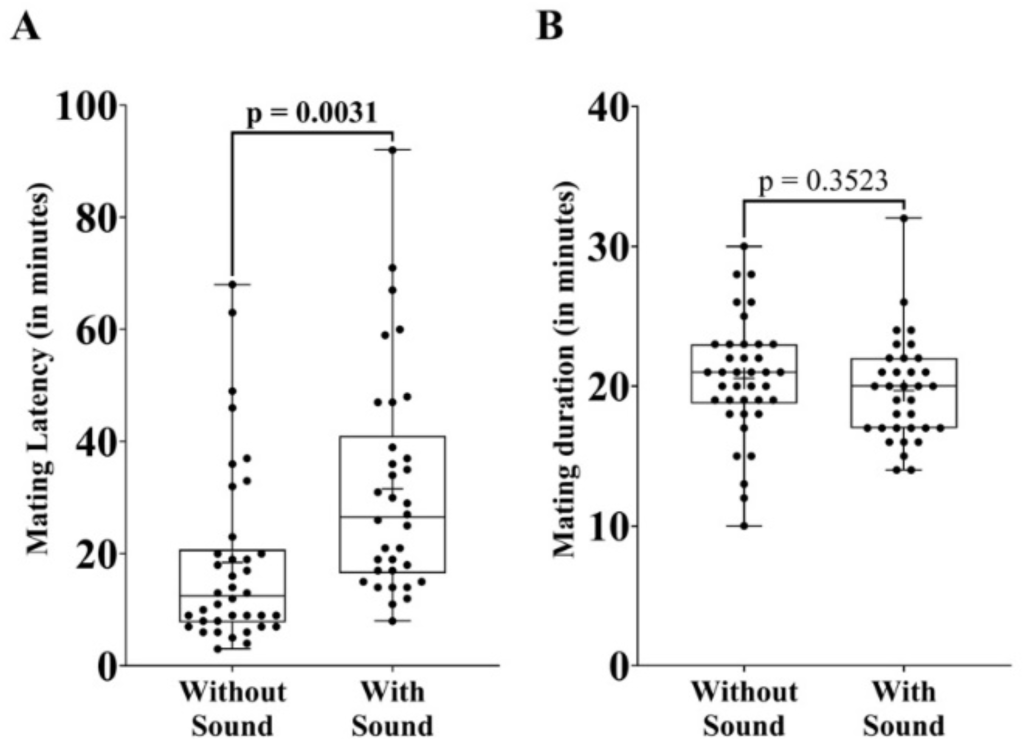
Effect of traffic noise on mating behaviour. A. Mating latency. B. Mating duration. n=40 pairs per treatment. Each pair consisted of a male and a female sampled independently from a large (∼2400 individuals) outbred lab-maintained population. Data points are shown by the jitter on the plot. The box’s top and bottom edges denote the 75^th^ and 25^th^ percentiles, respectively. The solid line between the edges represents the median. The whiskers denote the minimum and maximum data points. ‘+’ denotes the mean. Welch’s t-test (two-tailed) with a 95% confidence interval was performed to compare the means of the two treatments (i.e., with Sound and without Sound). The traffic noise exposure increased the mating latency of the flies but did not impact their mating duration.

### 3.3 Traffic noise exposure reduces locomotor activity in males, but females remain unaffected

We did a two-way ANOVA to test the effect of mating status and traffic noise exposure on locomotor activity. For the males, we fail to find a significant interaction between the mating status and traffic noise exposure (F_1_ = 1.85, *p* = 0.176). The main effect analysis suggested that mating status does not significantly affect the locomotor activity of males (F_1_ = 2.03, *p* = 0.157). The main effect analysis of traffic noise exposure showed a significant reduction in the locomotor activity of males when exposed to traffic noise (F_1_ = 19.96, *p* < 0.0001). For females, again, we do not find a significant interaction between the mating status and the traffic noise exposure (F_1_ = 1.203, *p* = 0.275). The main effects of mating status (F_1_ = 1.919, *p* = 0.169) and traffic noise exposure (F_1_ = 2.103, *p* = 0.15) on female locomotor activity were not statistically significant.

To check the effect of traffic noise exposure on the unmated and pre-mated flies separately, we performed Welch’s t-test and adjusted the p-values using the Holm Šídák procedure. We found that the response in the locomotor activity was sex-specific. In the case of locomotor activity of males, exposure to traffic noise led to a 34.3% reduction for the unmated (t_43.20_ = 2.163, p = 0.0361, d = 0.55, medium) (Fig 3A) and a 49.2% reduction for the pre-mated flies (t_52.43_ = 4.175, p = 0.0002, d = 1.062, large) (Fig 3B). For female locomotor activity, exposure to traffic noise did not lead to a significant change irrespective of flies being unmated (t_60.50_ = 0.2535, p = 0.8008, d = 0.064, small) (Fig. 3C) or pre-mated (t_55.24_ = 1.776, p = 0.0812, d = 0.444, small) (Fig. 3D).

**Figure 3:**
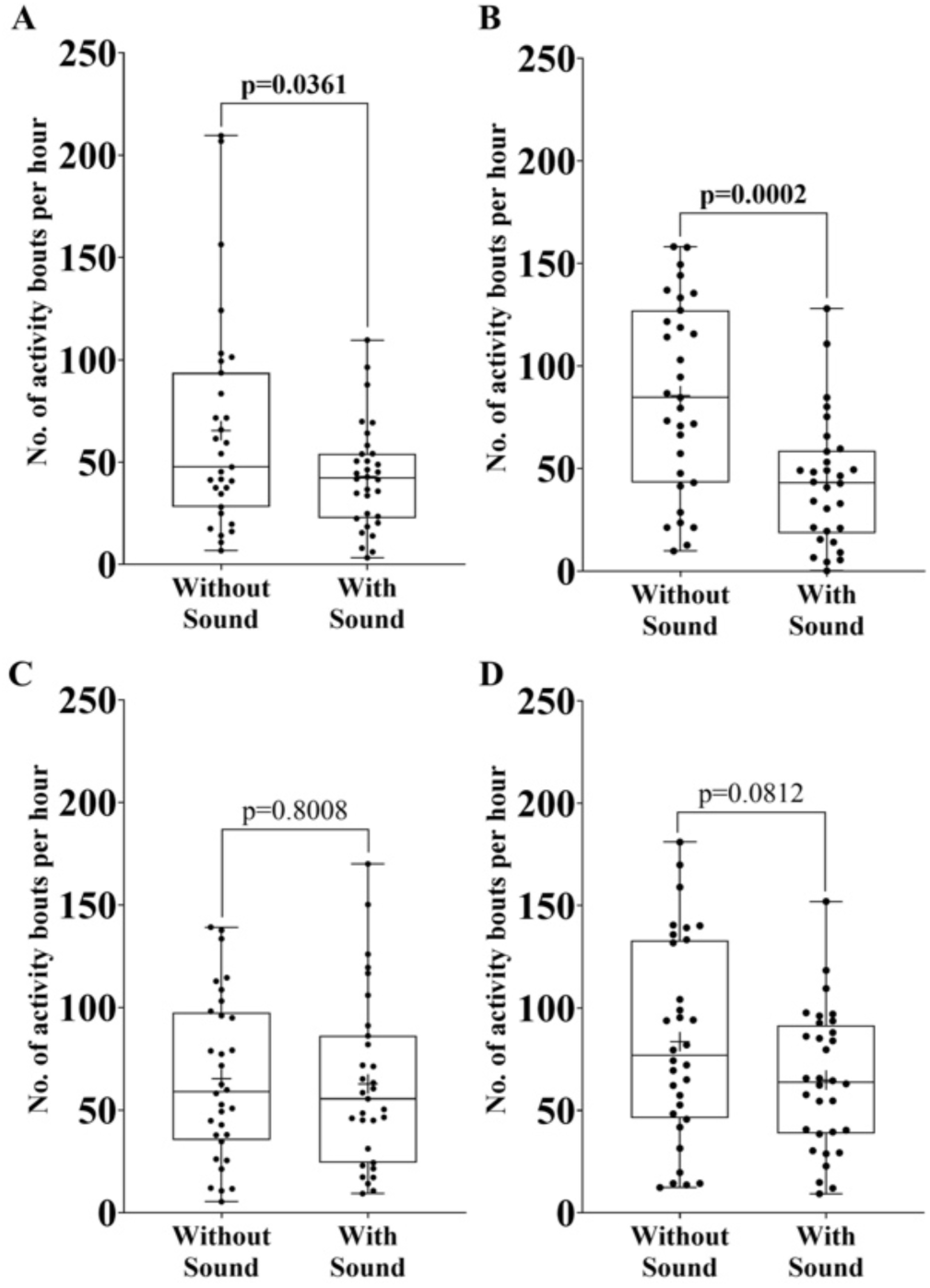
Effect of traffic noise on locomotor activity. A. Unmated males. B. Pre-mated males C. Unmated females D. Pre-mated females. n=32 per treatment. Each individual was sampled independently from a large (∼2400 individuals) outbred lab-maintained population. Data points are shown by the jitter on the plot. The box’s top and bottom edges denote 75^th^ and 25^th^ percentiles, respectively. The solid line between the edges represents the median. The whiskers denote the minimum and maximum data points. ‘+’ denotes the mean. Welch’s t-test (two-tailed) with a 95% confidence interval was performed to compare the means of the two treatments (i.e., with Sound and without Sound). P-values have been adjusted using the Holm-Sidak step-down procedure (for two comparisons). The results indicate that traffic noise exposure reduces the locomotor activity in males irrespective of the mating status, although there was no significant impact on the female locomotor activity.

### 3.4 Traffic noise exposure had no effects on desiccation resistance

In males, two-way ANOVA results reveal no significant interaction between the mating status (Unmated v/s Pre-mated) and the traffic noise exposure (With Sound vs Without sound) (F_1_ = 0.02, *p* = 0.89) on desiccation resistance. Main effect analysis suggested a significant reduction in the desiccation resistance of “pre-mated” male flies compared to the “unmated” male flies (F_1_ = 39.98, *p* < 0.0001) but no significant effect of traffic noise exposure on desiccation resistance (F_1_ = 2.08, *p* = 0.151). Again, we do not observe any interaction between mating status and traffic noise exposure in females (F_1_ = 0, *p* = 0.908). The main effect analysis for the female desiccation resistance reveals a significant increase in the desiccation resistance upon exposure to the traffic noise (F_1_ = 9.3, *p* = 0.003) and a significant reduction in the desiccation resistance of the mated female flies (F_1_ = 154.4, *p* < 0.0001).

No significant difference was observed between the desiccation resistance of the noise-exposed and the control flies, irrespective of them being unmated (t_61.73_ = 1.314, p = 0.1937, d = 0.33, small) (Fig. 4A) or pre-mated (t_37.83_ = 0.9109, p = 0.3681, d = 0.228, small) (Fig. 4B).

**Figure 4:**
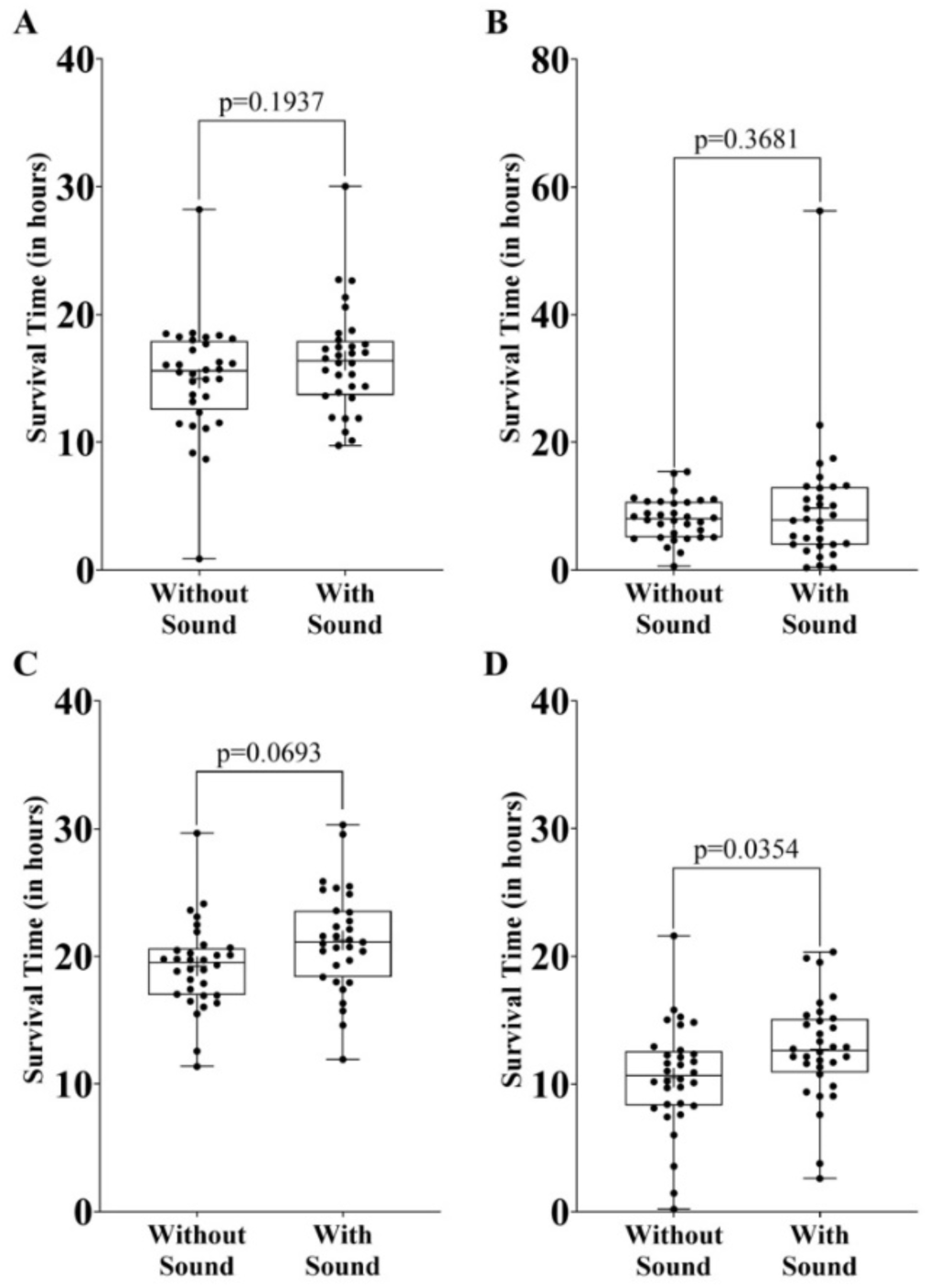
Effect of traffic noise on the desiccation resistance. A. Unmated males B. Pre-mated males C. Unmated females D. Pre-mated females. n=32 per treatment. Each individual was sampled independently from a large (∼2400 individuals) outbred lab-maintained population. Data points are shown by the jitter on the plot. The box’s bottom and top edges denote the 25^th^ and 75^th^ percentiles, respectively. The solid line between the edges represents the median. The whiskers denote the minimum and maximum data points. ‘+’ denotes the mean. Welch’s t-test (two-tailed) with a 95% confidence interval was performed to compare the means of the two treatments (i.e., with Sound and without Sound). P-values have been adjusted using the Holm-Sidak step-down procedure (for two comparisons). The results indicate that traffic noise does not have a statistically significant effect on the desiccation resistance of flies irrespective of their sex.

Welch’s t-test followed by Holm-Šídák correction show no significant change in the desiccation resistance of the female flies upon exposure to the traffic noise in both unmated (t_58.73_ = 2.155, p = 0.06935, d = 0.545, medium) (Fig 4C) and pre-mated flies (t_61.76_ = 2.151, p = 0.0354, d = 0.536, medium) (Fig 4D). Although the p-value for pre-mated female flies is less than 0.05, we treat it as non-significant following the conventions of the Holm-Šídák procedures, as a lower p-value had already been declared non-significant.

## 4 Discussion

In agreement with our initial hypothesis, the results from the study indicate that traffic noise exposure has significant adverse effects on the various life-history traits in *Drosophila melanogaster*. Traffic noise exposure reduced the reproductive output and locomotor activity (in males). Moreover, we observed the possibility of impaired communication between male and female mating pairs as the traffic noise exposure significantly increased the mating latency in the flies. These results indicate a clear fitness cost associated with living in environments continuously exposed to traffic noise.

Reproductive output (fecundity) is one of the most important life-history traits and could be instrumental in determining the organism’s fate in a habitat. We investigated the potential effect of traffic noise exposure on the fecundity of *Drosophila melanogaster*. Mating induces many physiological and behavioural changes in females, such as reduced receptivity to new males (Wolfner, 1997) and increased aggression (Bath et al., 2017). Therefore, to avoid possible confounds, we first assessed the fecundity of the flies without any previous mating experience (unmated). Freshly-mated flies exposed to traffic noise exhibited a large reduction in fecundity compared to the control flies (Fig. 1A). Upon repeating the assay twice on the independent sets of flies (freshly-mated) from the same population, we observed a similar significant reduction in fecundity as compared to the respective independent controls. Then we asked, what was responsible for this reduction in fecundity?

Mating has been shown to boost fecundity in *Drosophila* (Kalb et al., 1993; Wolfner, 1997), implying that any hindrance to mating could potentially reduce fecundity. In these flies, male courtship includes different behaviours like orientation, wing tapping, wing vibration, wing flicking, wing waving, wing semaphoring, wing scissoring, leg rubbing, leg vibration, circling, licking and mounting, followed by acceptance or rejection signals (involving less auditory and more mechanical components) from females (Spieth, 1974). Auditory courtship cues (’love song’ (Bennet-Clark and Ewing, 1970)), which are produced by the wing vibration in males, play an essential role in the reproductive success of *Drosophila* mating (Lasbleiz et al., 2006; von Schilcher, 1976), and masking of the love song due to the similar frequencies of the environmental noise could lead to mate rejection (Samarra et al., 2009). Due to its wide frequency profile (∼50 Hz to 10,000 Hz) (Can et al., 2010), the frequency components of traffic noise could lead to the masking of *Drosophila melanogaster* courtship songs (150 Hz to 300 Hz (von Schilcher, 1976)). These prior observations from the literature suggest that traffic noise exposure could hinder the mating behaviour of the flies. In spite of a significant increase in the mating latency of the flies when exposed to the traffic noise, the mating duration was not significantly affected. This suggests that traffic noise may have hindered only the courtship process, or the communication between the male-female pair before the male gets mounted on the female and once the stable mounting happens, the duration post that is unaffected. Another factor that might have played a part in increased mating latency, is the reduced male locomotor activity due to the traffic noise (as observed in the locomotor activity assay (Fig. 3A and 3B).

To test if the mating status of the flies drove the reduction in fecundity, we measured the number of eggs laid by flies that had 4 to 5 days to mate before the assay was conducted. If mating was affected due to exposure to the traffic noise and only the mating status was driving the reduction in the fecundity, we expected no significant reduction in the fecundity of the pre-mated flies upon exposure to the traffic noise. However, we observed a significant reduction in the fecundity of the pre-mated flies when exposed to the traffic noise compared to the flies that were not exposed to the traffic noise (Fig. 1B). Moreover, the magnitude of fecundity reduction was higher in freshly-mated flies (Fig. 1A) than pre-mated flies (Fig. 1B). Although our study does not allow us to understand the cause for this difference, some speculations can nevertheless be made. It is known that noise can induce oxidative stress in *Drosophila* (Dhar et al., 2020). Oxidative stress in mammals has been shown to reduce sperm count and quality (reviewed in Tremellen, 2008; Walke et al., 2023). The pre-mated females had mated in the absence of noise and therefore might have stored a large amount of good-quality sperm in their bodies. On the other hand, the freshly-mated flies had their first mating in the presence of noise, and hence did not have access to sufficient good-quality sperm, leading to a greater reduction in their fecundity than the controls. This speculation assumes that the effects of noise on sperm quality are similar in mammals and flies. Although plausible, we could not locate any study that has explicitly addressed this issue, thus providing a potentially interesting problem for future research.

The large decrease in fecundity and the increased mating latency (Fig. 2A) indicated a clear fitness cost associated with traffic noise exposure. We measured the reproductive output when the experimental flies were 13-day-old adults, for a window of 12 hours, during which the flies were continuously exposed to traffic noise. The lifespan of the outbred unselected *D. melanogaster* ranges from 60 to 90 days (Ziehm and Thornton, 2013; Ziehm et al., 2013). It is suggested in the literature that the net reproductive output over the individual’s lifetime is a better measure of the individual’s fitness (Krimbas, 2004). Also, organisms with lower reproductive output in early life could have a longer lifespan, leading to a longer reproductive phase in the life cycle, eventually compensating for the low fecundity (Djawdan et al., 1996). However, depending upon the magnitude and nature of the stress and the extent to which fecundity has been reduced, it might often not be possible for the lifespan to increase to the extent to compensate for the reduced fecundity fully; the worst case scenario could be that the lifespan and fecundity decrease together. Indeed, exposure to acoustic stimuli decreases the lifespan in *D. melanogaster* males (although not in females) (Morales et al., 2010). Noise has been reported to induce increased physiological oxidative stress and the possibility of organ damage in *Drosophila* (Christie et al., 2013). In the real world, we expect the flies to experience traffic noise throughout their lifespan with no relief from the stress imposed due to it. Therefore, the decrease in fecundity in our study will most likely translate into a lifetime decrease in the reproductive output in the real world.

Acoustic trauma could lead to behavioural changes like increased anti-social behaviour in rats (Zheng et al., 2011) and reduced aggressive behaviour in fruit flies (Dhar et al., 2020). Locomotor activity is at the center of almost all kinds of behaviours that require some movement, be it aggression, foraging (Van Alphen and Jervis, 1996), courtship (Greenspan and Ferveur, 2000), exploration (Bell, 1990) and circadian rhythm (Dunlap, 1999). Locomotor activity also influences predation escape behaviour (Card, 2012). Therefore, locomotor activity plays a vital role in determining the fitness of the organisms. We investigated the effect of traffic noise exposure on the locomotor activity of *D. melanogaster*. Irrespective of their mating status, we observed a decrease in male locomotor activity (decreased activity bouts per unit time) when exposed to traffic noise (Figs 3A and 3B). On the other hand, the female locomotor activity was not significantly affected by the exposure to traffic noise (Figs 3C and 3D). This reduced activity of the male flies, combined with the adverse effects on the other life history traits, could further reduce the fitness of the individuals.

Due to their small body sizes, the surface area to volume ratio of insects is quite large, making them highly susceptible to desiccation stress (Gibbs, 2002). Vehicular emissions contribute significantly to increasing the rate of global warming (Alam and Khan, 2020). Climate change and global warming further deteriorate the conditions by increasing the risk of desiccation. Moreover, mating is known to increase the desiccation resistance in some species of *Drosophila* (Knowles et al., 2004). On the other hand, since fecundity goes down due to noise exposure, there is a possibility that such flies have a greater amount of resources left to combat stresses like desiccation. Thus, we sought to confirm whether there was a change in the desiccation resistance of the flies exposed to noise. We did not observe any significant change in the desiccation resistance of flies (both males and females) when exposed to the traffic noise (Fig. 4). The desiccation resistance in the pre-mated flies was significantly lower than that of the unmated flies in both sexes, which might result from post-mating physiological costs.

We note here that we have used only one sound clip for the noise treatment in this study. Therefore, in principle, the effects we observed could be due to some features of the specific sound clip, rather than traffic noise in general. The sound profile of the used clip was very much within the known frequency and intensity ranges of traffic noise (Can et al., 2010; Grubesa and Suhanek, 2020). Moreover, we did not observe any peculiarities in our sound clip (e.g., only one kind of vehicular noise). Therefore, we believe our results should apply to traffic noise in general. However, a conclusive demonstration of generalizability demands that our entire study be repeated under multiple independent instances of traffic noise. While out of the scope of the present study due to logistical constraints, such a demonstration of generalizability would certainly add value to the field.

## 5 Conclusion

Noise can negatively affect the fitness of insects like crickets (Gurule-Small and Tinghitella, 2019; Welsh et al., 2023) and ants (Cammaerts and Cammaerts, 2018). Our study extends this line of work into traffic noise, which has certain specific properties (heterogeneous and broader range of intensity and frequency profile) and has become a significant source of pollution in the real world. We found that exposure to traffic noise can affect multiple life-history parameters in the fruit fly, *Drosophila melanogaster*, which can potentially reduce their fitness. This implies that the ever-expanding road networks can adversely affect insects and the entire ecosystem. However, we note here that the overall effects of noise on arthropod diversity are complex, and some taxa can also benefit or remain unaffected by anthropogenic noise (Bunkley et al., 2017). Therefore, much more data is needed to understand the effects of anthropogenic noise on insects, particularly under natural conditions.

## Acknowledgements

We acknowledge Hareendran KM and Navanath Kadam for helping with the experiments, as well as Akshay Malwade and B. Vibishan for helpful discussions.

## Data Availability

All data will be deposited into an open-access database upon acceptance.

## Supplementary Information

**Figure S1:**
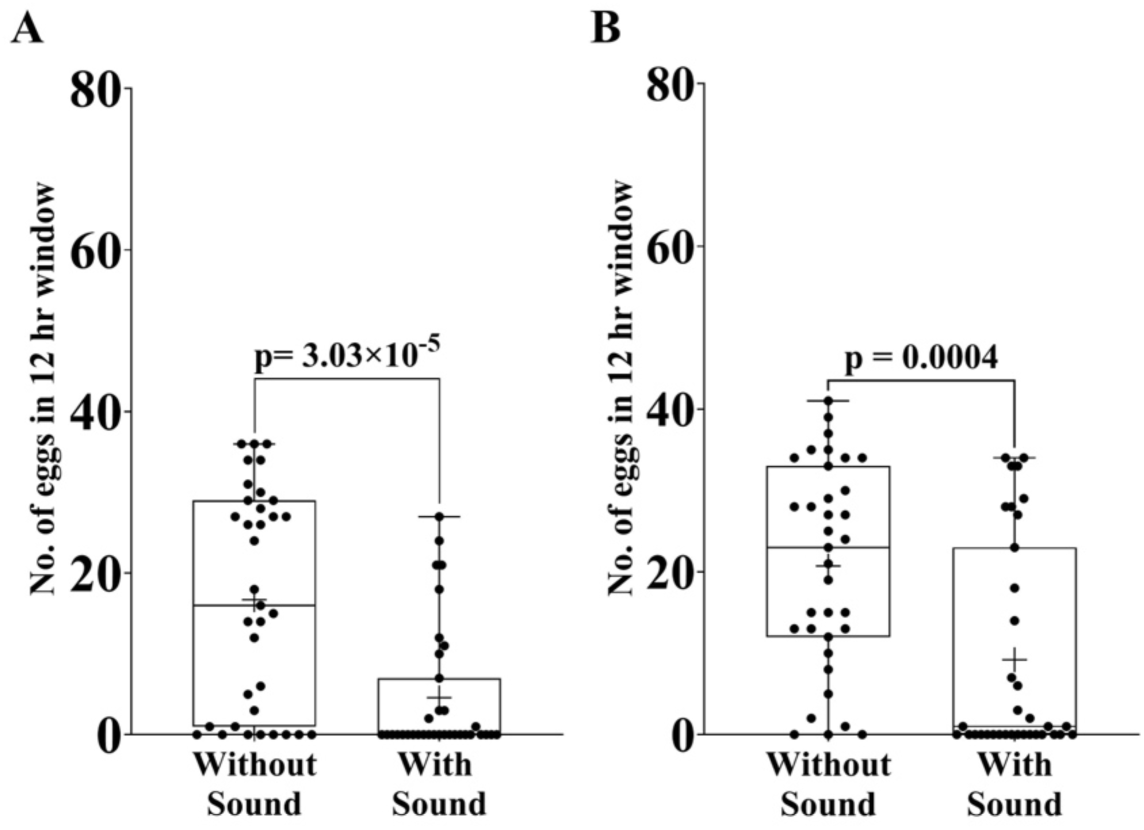
Repeat experiment data for the effect of traffic noise on fecundity. A. Freshly-mated (repeat 1), B. Freshly-mated (repeat 2). n=35 pairs for per treatment per experiment. Each pair consisted of a male and a female sampled independently from a large (∼2400 individuals) outbred lab-maintained population. Data points are shown by the jitter on the plot. The top and bottom edges of the box denote the 25^th^ percentile and the 75^th^ percentile, respectively. The solid line between the edges represents the median. The whiskers denote the minimum and maximum data points. ‘+’ denotes the mean. Welch’s t-test (two-tailed) with a 95% confidence interval was performed to compare the means of the two treatments (i.e., with Sound and without Sound). P-values have been adjusted using the Holm-Šídák step-down procedure (for two comparisons).

**Figure S2:**
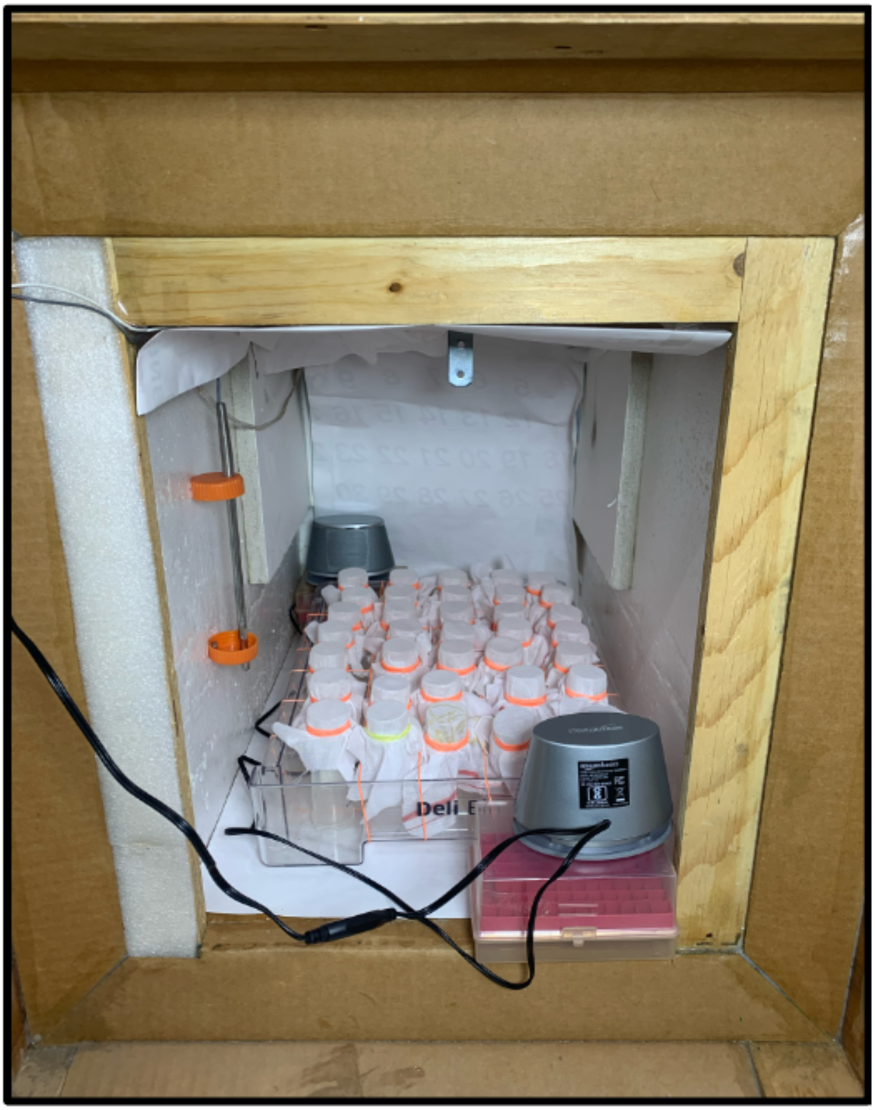
Wooden Chamber with fecundity setup. The door to the chamber is kept open so one can click this picture.

**Figure S3:**
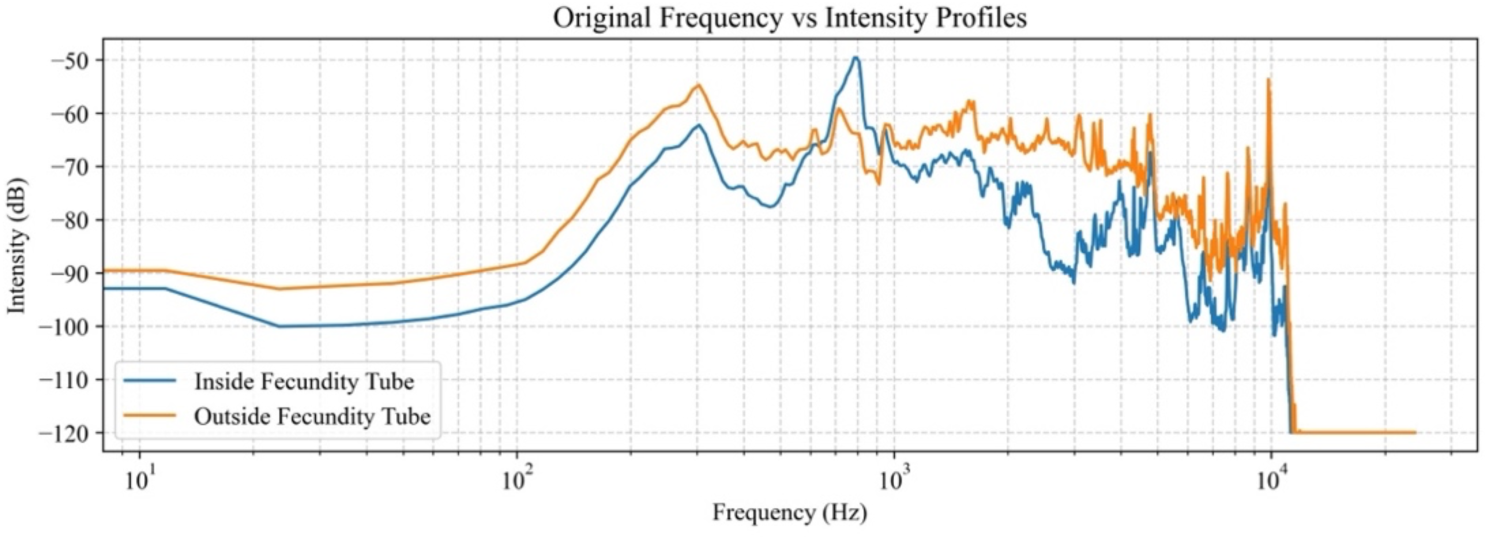
Comparison of the spectrum of the noise playback clip inside and outside the fecundity tube. We recorded the traffic noise playback outside and inside the fecundity tube and compared the frequency and intensity profiles using scipy.signal.welch in SciPy package in Python. The results indicate a similar frequency profile between the spectrum inside and outside the fecundity tube with some fluctuations at the higher frequencies.

**Figure S4:**
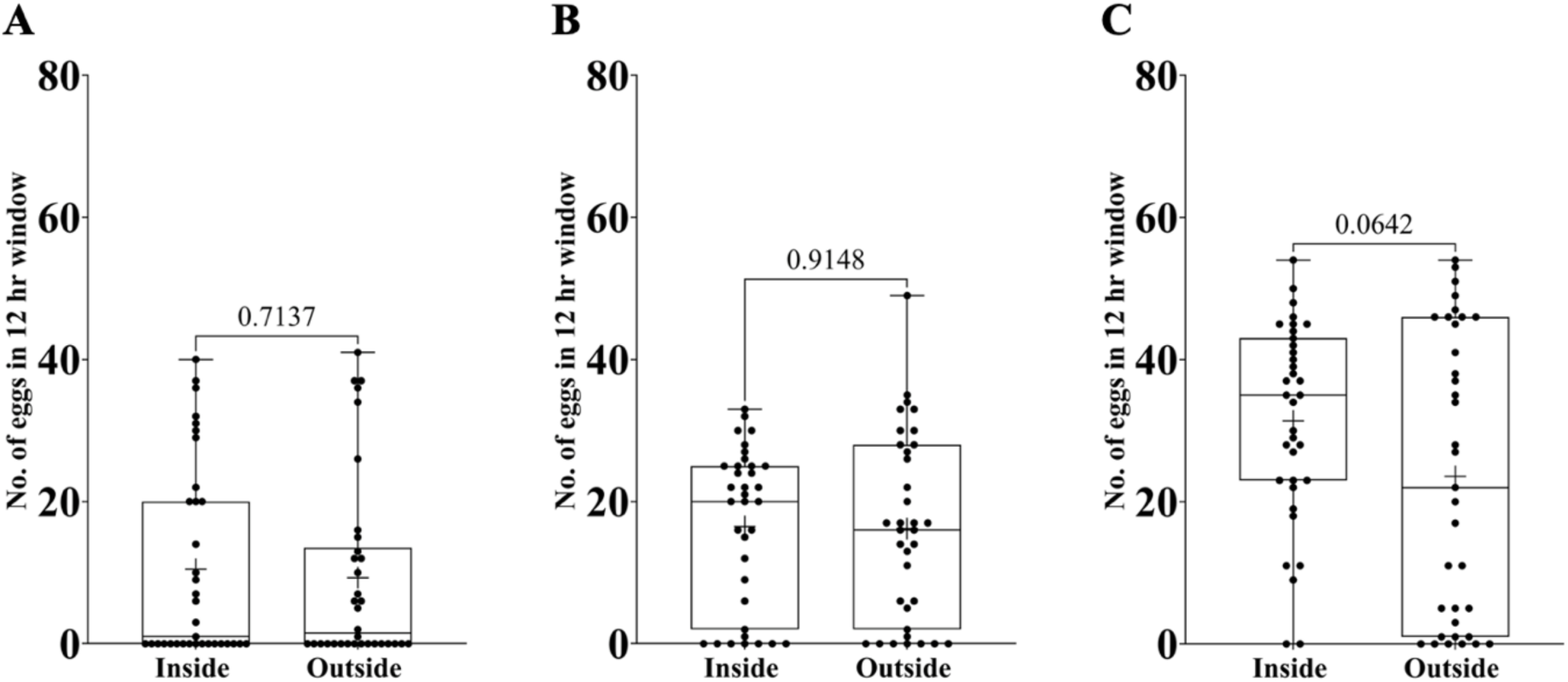
Control Experiments to check for the effect of lighting and environment inside and outside the wooden chamber on the fecundity of the flies (in the absence of traffic noise exposure) A. Control Experiment 1 B. Control Experiment 2 C. Control Experiment 3. n=35 pairs for per treatment per experiment. Each pair consisted of a male and a female sampled independently from a large (∼2400 individuals) outbred lab-maintained population. Data points are shown by the jitter on the plot. The top and bottom edges of the box denote the 25^th^ percentile and the 75^th^ percentile, respectively. The solid line between the edges represents the median. The whiskers denote the minimum and maximum data points. ‘+’ denotes the mean. P-values shown were obtained from Welch’s t-test (two-tailed) with a 95% confidence interval, which was performed to compare the means of the two treatments (i.e. Inside and outside the wooden chamber).

